# Natural language processing of gene descriptions for overrepresentation analysis with GeneTEA

**DOI:** 10.1101/2025.03.28.646026

**Authors:** Isabella A Boyle, Nayeem Akram Aquib, Mustafa Kocak, Randy Creasi, Phil G Montgomery, Catarina D Campbell, Joshua M Dempster

## Abstract

Overrepresentation analysis (ORA) is used to identify the biological relationships in a list of genes by testing gene sets for enrichment in the query. However, the inconsistent definition and highly overlapping nature of gene set databases can make interpreting ORA results difficult. Here, we introduce GeneTEA, a model that takes in free-text gene descriptions and incorporates several natural language processing methods to learn a sparse gene-by-term embedding, which can be treated as a *de novo* gene set database. We benchmark performance against other popular ORA tools and find that only GeneTEA properly controls false discovery while consistently surfacing relevant biology.

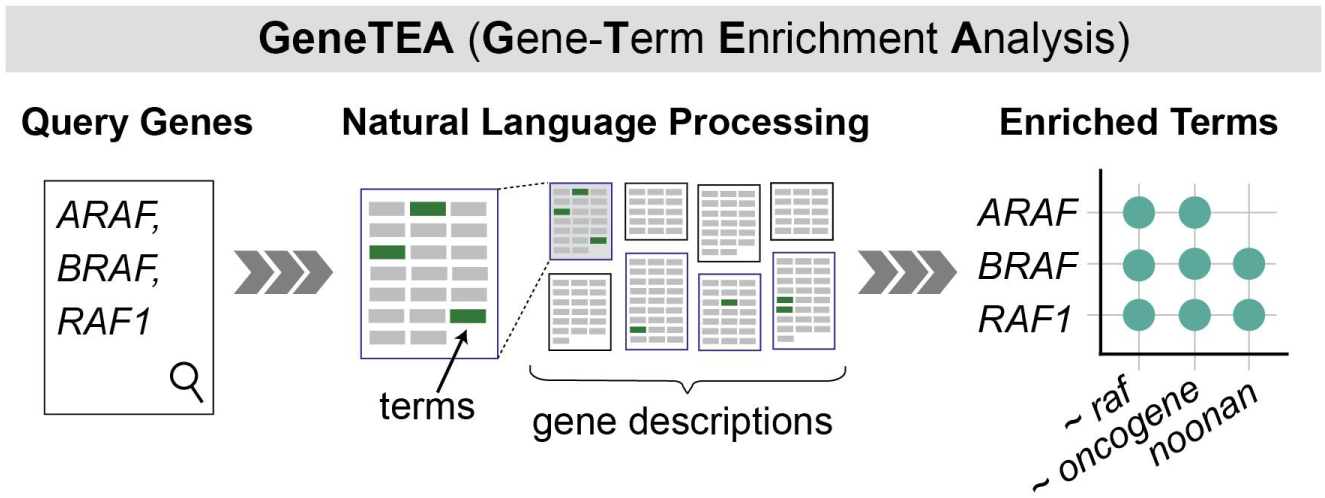

## INTRODUCTION

Technological advances have enabled a shift toward genome-scale, hypothesis-generating experiments. Accordingly, overrepresentation analysis (ORA) was developed to gain biological insights into these high-dimensional data. This method involves testing a query list of genes derived from large-scale experiments for statistical enrichment of gene sets encoding biological processes, molecular functions, phenotypes, or other pre-existing knowledge (1). There is no universal definition of a gene set, and today they are defined orthogonally across many databases, including the Gene Ontology (2), Human Phenotype Ontology (3), Molecular Signatures Database (4), Kyoto Encyclopedia of Genes and Genomes (KEGG) (5), WikiPathways (6) and Reactome Database (7).

Many tools have been developed to run ORA across these databases simultaneously, with g:GOSt (8) and Enrichr (9) among the most popular. However, these models are only as informative as the gene sets they test on - and the explosion in gene set databases has resulted in many redundant, conflicting, and poorly defined gene sets. Additionally, the highly overlapping nature of the gene sets within and between these databases has been shown to reduce the specificity of ORA (10). Furthermore, the magnitude of significance values is directly related to the size of the gene set database queried, making it difficult to interpret results from tools that aggregate results across many databases (11).

In recent years the field of natural language processing (NLP) has experienced many impressive advancements in text representation. Among the simplest NLP models are bag of words (BoW) models, which treat text as a countable collection of unordered tokens (words or phrases); while large language models (LLMs) learn context-dependent embeddings influenced by the position of tokens (12). Some have proposed LLMs could be leveraged to identify relationships via prompt engineering (13); for example, TALISMAN (14) and GeneAgent (15) provide tailored prompts for generating names and summaries for lists of genes. However, LLMs are incapable of reporting statistics and are therefore unable to quantify if the relationships described by their outputs are statistically significant. Furthermore, post hoc steps must be taken to verify the generative output is not the result of hallucination, a phenomenon where LLMs produce seemingly coherent but factually incorrect outputs. These verification processes measure the semantic similarity of the generated name to the results of ORA on existing gene set databases, effectively restricting these prompting strategies to reporting enrichments that traditional ORA could identify.

A recent comparative study of biomedical reasoning NLP tasks found that LLM performance is highly sensitive to prompt engineering and achieved equivalent results to a simpler BoW model (16). A potential explanation for this is that LLMs trained on massive corpora scraped from the internet encounter many incomplete, outdated, and conflicting research results (17). For example, one group performed text-mining on a corpus of more than 200, 000 abstracts from dementia publications and found that 91% (325/335) of KEGG pathways were implicated in Alzheimer’s Disease by 5 or more studies (18).

We hypothesized that identifying enriched terms in gene descriptions would capture equivalent information to what is encoded in gene set databases, and do so without requiring a priori defined gene sets - thus reducing redundancy and increasing interpretability. A similar idea explored by Leong and Kipling (19), which tested whether single-word tokens extracted from PubMed papers could be used for ORA, showed promise. However, this approach preceded many advancements in NLP - leaving room for improvement in surfacing relevant terms, eliminating stopwords (common, uninteresting words), and consolidating synonymous tokens. Furthermore, to ensure the corpus concretely linked text to specific genes, we opted to utilize gene descriptions as these texts distill general knowledge about protein function, localization, and pathway involvement alongside major findings from the literature. A gene-by-term embedding learned by a BoW model could then be treated as a unified gene set database for ORA, and LLMs could be leveraged to identify stopwords and synonymous terms. Incorporating these concepts, we developed the Gene-Term Enrichment Analysis (GeneTEA) model (Figure 1). To assess GeneTEA, we analyzed its gene-by-term embedding, benchmarked its performance against competitor ORA tools, and tested the framework’s flexibility in generalizing to other organisms’ genomes and other biological entities such as drugs.

**Figure 1.**
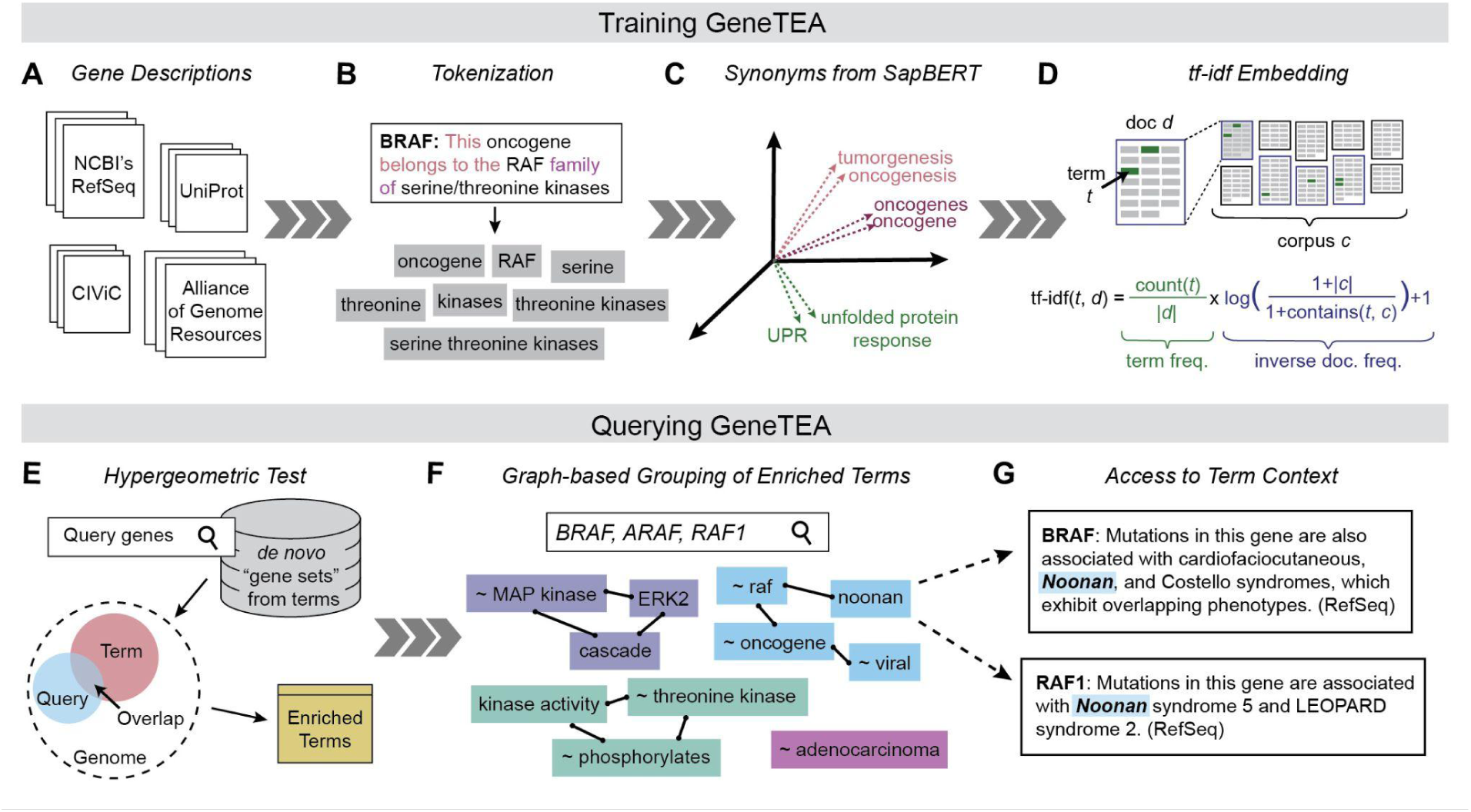
Overview of the GeneTEA model. (**A**) The training corpus was constructed from free-text gene descriptions. (**B**) Example of tokenization. (**C**) Example SapBERT embeddings for tokens, colored by the assigned synonym set. (**D**) Diagram and equation representing term frequency-inverse document frequency (*tf-idf*) embedding. (**E**) Graphic representing the use of a hypergeometric test to identify enriched terms. (**F**) Example of the term groups for the query BRAF, ARAF, RAF1. (**G**) Text excerpts referencing the term “noonan” in BRAF and RAF1.

## MATERIAL AND METHODS

### Design of the GeneTEA model

The GeneTEA model was constructed around a corpus of free-text gene descriptions from public resources including NCBI’s RefSeq (20), UniProt (21), Clinical Interpretation of Variants in Cancer (CIViC) (22), and the Alliance of Genome Resource’s natural language summary of the Gene Ontology terms (23) (Figure 1A, Supplementary Note S1). Each text was split into sentences which were then tokenized into words. Additionally, phrases defined in the UMLS Metathesaurus (24) were extracted, to retain only biologically meaningful *n*-grams (Figure 1B, Supplementary Note S2).

We noticed many synonymous terms were present in the vocabulary - such as “oncogene” and “oncogenes” or “unfolded protein response” and “UPR” - and sought to identify them systematically using semantic similarity. Therefore, we clustered token embeddings from SapBERT (25), an LLM fine-tuned on PubMed, using HDBSCAN (26) over pairwise cosine distance (Figure 1C, Supplementary Note S3). We compared this unsupervised synonym set assignment to lists of manually curated concept names specified in UMLS Metathesaurus and found strong agreement between the clusterings (Figure S1A). These learned synonym sets were labeled by the most frequent term with the prefix “∼” and treated as drop-in replacements for all terms in the cluster.

After texts were preprocessed, a term frequency-inverse document frequency (*tf-idf*) embedding was constructed, such that gene description documents were represented as a gene-by-term sparse matrix (Figure 1D, Supplementary Note S4). The *tf-idf* embedding is a popular BoW representation in NLP, which scales the number of appearances of a term in a document (term frequency, *tf*) by the number of documents containing that term across the corpus (inverse document frequency, *idf*). This normalization up-weights rarity and repetition, attributing higher values to more salient terms. In fact, *tf-idf* values can be interpreted as the mutual information between a term and a document (27). This gene-by-term matrix could then be binarized to produce a *de novo* gene set database learned from the corpus of gene descriptions.

When querying GeneTEA, the user provides a list of genes. Per term, the number of genes in the query whose description contains that term is counted and a hypergeometric test is performed to determine overrepresentation (Figure 1E, Supplementary Note S5). This *p*-value is corrected with Benjamini-Hochberg’s method for multi-hypothesis testing to produce a false discovery rate (FDR) value. An FDR threshold of 0.05 is applied to obtain enriched terms and the sum of *tf-idf* values across the query is reported as the effect size, which determines the sort order. Any stopwords (Supplementary Note S6) or terms matching only a single gene are filtered out. Additionally, strict sub-terms are removed if they are less informative than a longer super-phrase (Supplementary Note S7).

For queries with strong underlying biological signals, many related - but not strictly synonymous - terms may arise. To ease interpretation, we employ a graph-based grouping strategy that condenses terms with associated meanings in the context of the query (Figure 1F, Supplementary Note S7). Concretely, a graph connecting terms by their similarity in the *tf-idf* embedding is used to identify communities of conceptually related terms. To account for context specificity, the edge weight is the product of the cosine similarity of a pair of terms within the query and across the entire corpus. Low weight edges are pruned, then a greedy-modularity algorithm is used to identify communities. For example, in a query with BRAF, ARAF, and RAF1 the terms “∼ raf”, “∼ oncogene”, “noonan”, and “∼ viral” form a group since in this context these describe the RAF families’ initial discovery as a viral oncogene in murine models. These groups are then labeled by the three most informative terms - for example, the example above would be labeled “∼raf | ∼ oncogene | noonan”.

A critical advantage of GeneTEA’s construction is that the link between the term and source text is retained, allowing a user to refer directly to the context in which that term was used (Figure 1G). For example, “noonan” could be a puzzling term to encounter, but by reviewing the text excerpts from the query genes’ descriptions it becomes apparent this term refers to Noonan Syndrome, a disease associated with germline mutations in the RAF and RAS families (28). In addition to enabling users to assess the reliability of the enrichment, this functionality eases the process of hypothesis building by establishing a direct explanation for how query genes relate to the terms identified.

### Validation and Benchmarking

We performed various analyses to validate GeneTEA and benchmark its performance against other ORA tools. The following sections define these procedures in more detail.

#### Latent Semantic Analysis

To better understand the underlying biological signal in GeneTEA’s embedding, we performed latent semantic analysis (LSA) using sklearn (v1.4.0) (29). First, the *tf-idf* matrix was reduced to 500 dimensions using TruncatedSVD decomposition with ARPACK solver. Then, the matrix was normalized using Normalizer. Finally, pairwise cosine similarities were computed along the gene axis.

#### Gene set overlaps

To assess the overlap between gene sets within a database, we computed the probability of two randomly selected gene sets having high overlap. Specifically, if the smaller gene set in the pair had at least half its members in common with the larger gene set it was considered highly overlapping. We estimated the probability of high overlap by randomly sampling 500 gene sets from each of the gene set databases underlying GeneTEA, g:GOSt, and Enrichr and found the percent of pairs with high overlap. This process is repeated 10 times to assess stability.

#### Querying competitor ORA models

For g:GOSt, we obtained human gene set enrichment results from all available libraries via the gprofiler python package’s “profile” function (v1.0.0). To filter to the top terms, we retained only entries matching at least 2 query genes annotated as “significant” as determined by the provided adjusted *p*-value threshold. Significance defined the sort order. In cases where the same term was reported in the output multiple times, we retained only the row corresponding to the first appearance.

For Enrichr, we queried the “enrich” API endpoint (on 2025-03-27) for each tested library and concatenated the results. We used the 130 libraries presented on the Enrichr website’s “Transcription”, “Pathways”, “Ontologies”, “Diseases/Drugs”, and “Cell Types” tabs as of 2025-03-03. We retained only entries matching at least 2 query genes where the “Adjusted p-value” was below 0.05 and sorted by “Combined score.” In cases where the same term was reported in the output multiple times, we retained only the row corresponding to the first appearance.

#### Assessment of false discovery control

We created a benchmarking set of random queries to assess false discovery control. We randomly sampled lists of 3, 5, 10, 15, 20, 25, 50, 100, 250, 500, and 1000 genes 10 times each, resulting in 110 queries. We then counted the number of enriched terms identified by each model for each query, representing false discoveries.

#### MedCPT Relevance and Redundancy metrics

To quantify the relevance and redundancy in a model’s enriched terms, we turned to MedCPT (30). We utilized the huggingface API (transformers v4.46.3) to access NCBI’s MedCPT CrossEnc (“ncbi/MedCPT-Cross-Encoder”) and AEnc (“ncbi/MedCPT-Article-Encoder”). For each tested gene set, we treated the comma-separated list of genes as the query and either a single term or the period-separated top 100 enriched terms from each model as the set of documents. To measure MedCPT Relevance, we embedded the pairs of query and document with the CrossEnc’s tokenizer, then fed this to the model to obtain the output logits. We took the top 100 enriched terms from each model, embedded them with the AEnc, and counted the number of pairs with cosine similarity >0.95 to quantify redundancy.

#### Query gene sets used for benchmarking

To create random samples of the Hallmark Collection (31), we sampled lists of genes of length 10, 25, 50, and 100 from each of the 50 gene sets to create 200 queries. We then randomly combined pairs of these sampled sets to create an additional 100 queries.

To create a set of experimentally derived queries, we obtained lists of genes with length >2 from the following sources:

- “AlphaFold2 Protein Clusters”: Data file “3-sapId_sapGO_repId_cluFlag_LCAtaxId.tsv.gz” from Barrio-Hernandez et al. 2023 (32) downloaded from https://afdb-cluster.steineggerlab.workers.dev/. UniProt IDs were mapped to gene symbols using the HUGO Gene Nomenclature Committee (HGNC) mapping (33), and only clusters representing >2 HGNC gene groups were retained.
- “BioID Interacting Proteins”: Supplementary Table 4 of Go et al. 2021 (34)
- “Human Disease GWAS with Rare Variants”: Supplementary Table 8 of Wang et al 2021 (35)
- “Perturb-seq Expression Modules” and “Perturb-seq Perturbation Clusters”: Supplementary Table 3 of Replogle el al. 2022 (36)

## RESULTS

### GeneTEA’s embedding efficiently encodes biological prior knowledge

To examine the biological signal captured by GeneTEA, we performed latent semantic analysis (LSA) on protein-coding genes to reduce the dimensionality to 500 components. We then evaluated gene-gene similarity by calculating the pairwise cosine similarity over the LSA embedding (Figure 2A). We observed that gene groups defined by HGNC, for example, the Histamine receptor family of genes, had high average cosine similarity (Figure 2B-C), which demonstrated that genes defined as related by an orthogonal source were represented by similar term embeddings in GeneTEA.

**Figure 2.**
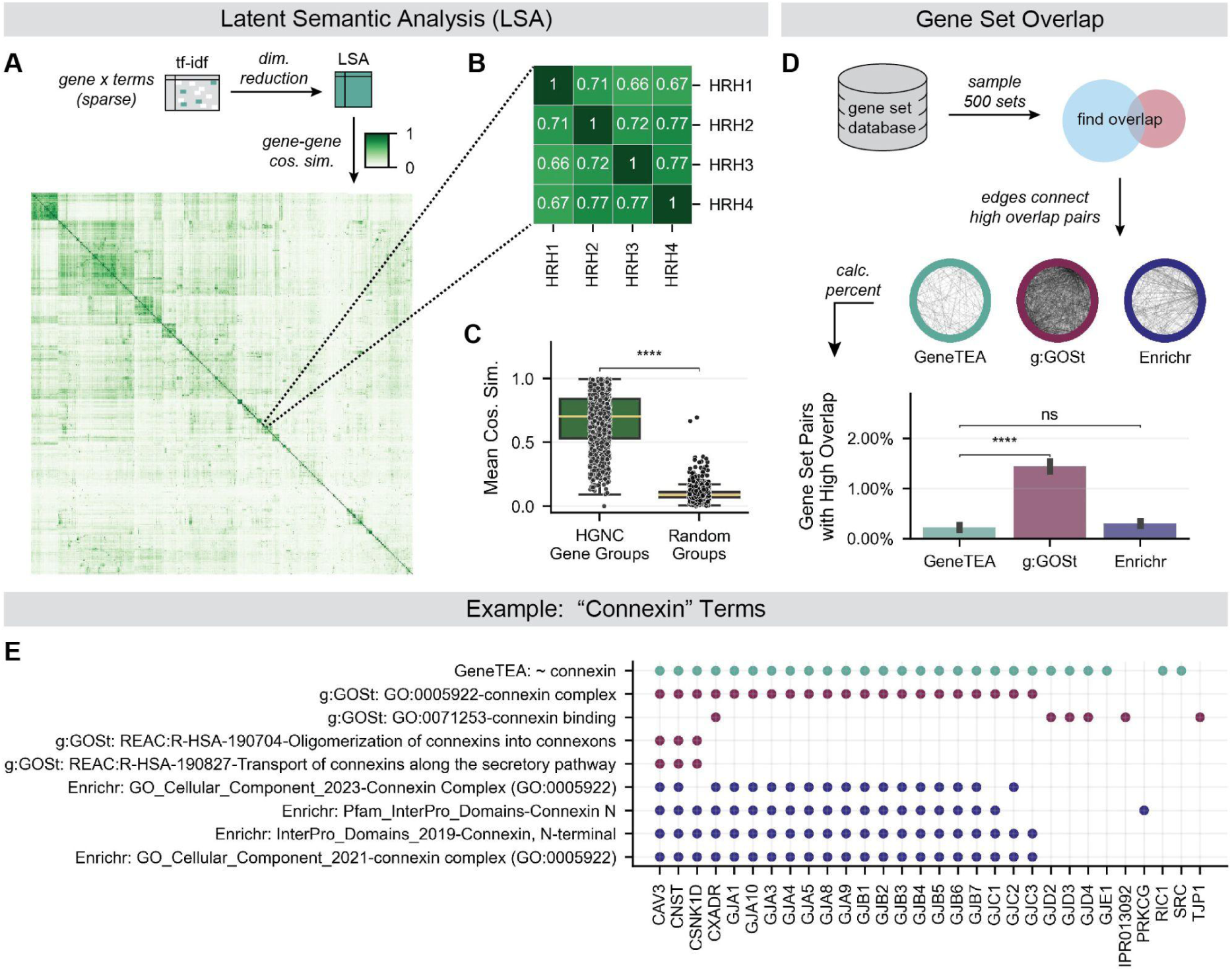
Examination of GeneTEA’s underlying embedding. (**A**) Upper: Latent semantic analysis (LSA) embedding was produced via a dimensionality reduction GeneTEA’s *tf-idf* matrix. Lower: Clustermap of cosine similarity between protein-coding genes in HGNC groups with <5 members (*n*=1,689) in LSA embedding. (**B**) Cosine similarity of HGNC’s Histamine receptor gene group in LSA embedding. (**C**) Average cosine similarity for HGNC’s gene groups in LSA embedding, compared to an equal number of random groups of equivalent size (*n*=1,443). (**D**) Upper: Hairball plots where nodes are a sample of 500 gene sets from each model’s database and edges represent a high overlap between a pair of gene sets. Lower: Bar plot comparing the mean percent of gene set pairs with high overlap across 10 samples, with error bars representing the standard error of the mean. (**E**) Example of gene sets from each database related to connexins, where points indicate whether a gene on the x-axis belongs to the gene set on the y-axis. Significance values are from a right-tailed Student’s *t*-test in **C** and a left-tailed Student’s *t*-test in **B**, where **** *p*-value < 0.0001 and ns indicates not significant.

GeneTEA’s embedding contains ∼24k terms (Figure S1B), which is nearly half the size of g:GOSt’s database (∼44k terms) and an order of magnitude smaller than Enrichr’s database (∼226k terms). Despite its compactness, we found that only a small percentage of pairs of gene sets had high overlap in GeneTEA (Figure 2D). This highlights that conflicting definitions from combining different ontology/pathway resources can lead to many highly overlapping gene sets - a problem GeneTEA avoids as it constructs gene sets from free-text. For example, in GeneTEA’s database, a single term identifies genes related to connexins, while in g:GOSt and Enrichr each contain 4 gene sets - various Gene Ontology terms, Reactome pathways, and InterPro domain-containing proteins (Figure 2E).

### GeneTEA yields reliable and relevant ORA results

We compared GeneTEA head-to-head with g:GOSt and Enrichr to benchmark our ORA model on several tasks. These competitor models also utilize a hypergeometric test; however, they apply differing multiple hypothesis correction strategies: Enrichr uses Benjamini-Hochberg FDR correction similar to GeneTEA, while g:GOSt employs a simulation-based *p*-value adjustment strategy to limit false discovery to 5%. For a fair comparison, we applied the equivalent FDR threshold of 0.05 to Enrichr and GeneTEA.

To test how well GeneTEA and its competitors control false discovery, we queried the models with 110 random gene sets of various lengths between 3-1000 and recorded the number of enriched terms (Figure 3A). GeneTEA produced false discoveries in only 1 of the random queries tested (<1%). In contrast, g:GOSt and Enrichr produced false discoveries in 46.4% and 69.1% of random queries, respectively. In total, Enrichr produced 1051 false discoveries while GeneTEA produced only 1, and g:GOSt yielded 116 (Figure S1C). Enrichr’s false discovery control failed more often in shorter length queries (<100 genes) while g:GOSt showed the inverse pattern and tended to produce more false discoveries as query length increased (Figure S1C-D). Together, these results indicate that GeneTEA is less susceptible to false positives than competitors, a crucial characteristic when generating hypotheses from ORA results.

**Figure 3.**
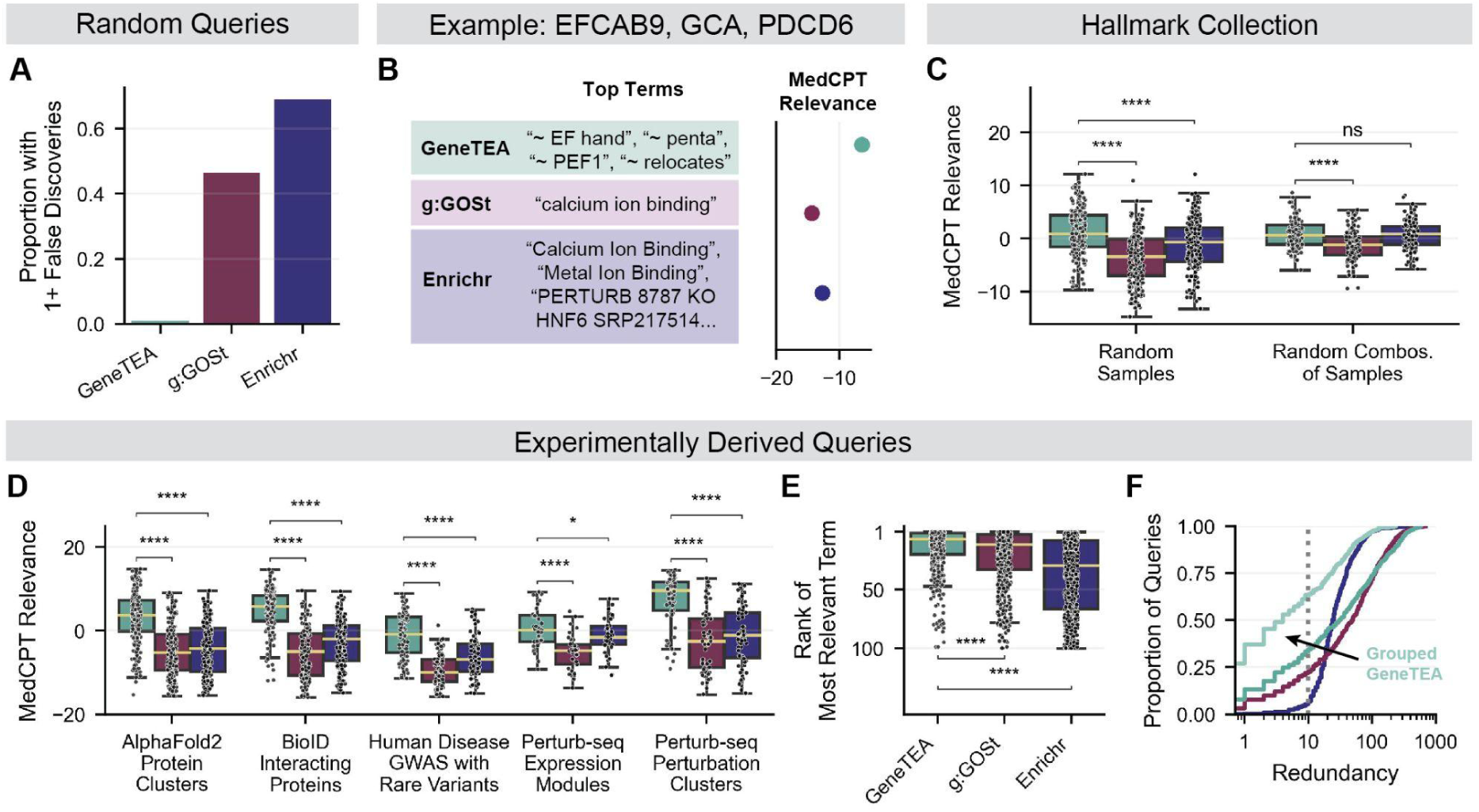
Performance of GeneTEA and competitors on a variety of benchmarking tasks. (**A**) The proportion of random queries (*n*=110) that resulted in 1 or more false discoveries. (**B**) Example of MedCPT Relevance metric. For genes encoding three EF-hand proteins, the top 100 terms from each model are obtained and ranked by MedCPT, resulting in a unitless continuous value where a more positive value indicates higher relevance. (**C**) MedCPT Relevance for random samples of gene sets in the Hallmark Collection (*n*=200) and random combinations of these samples (*n*=100). (**D**) MedCPT Relevance for various experimentally derived queries (*n*=532). (**E**) Rank of the most relevant term in a model’s enrichment results for a query, as indicated by highest MedCPT Relevance. (**F**) Cumulative distribution plot of redundancy (semantically similar term pairs) across experimentally derived queries (*n*=532). All significance values in **C**-**E** are from a right-tailed Student’s *t*-test, where * *p*-value < 0.05, **** *p*-value < 0.0001, and ns indicates not significant.

To assess an ORA model’s ability to detect biological signals, we needed a method for measuring the relevance and redundancy of enriched terms. For this purpose, we utilized MedCPT (30), a biomedical information model built to perform lexical matching for semantic retrieval based on 255 million user click logs from PubMed. As MedCPT ranks free-text articles against a user’s search query, it produces a continuous, unitless score for each article where higher values indicate higher relevance for a given search. Therefore, we report this score for the top terms from each model as the MedCPT Relevance (Figure 3B). For example, GeneTEA’s enriched terms (“∼ EF hand”, “∼ penta”, “∼ PEF1”, “∼ relocates”) for the query *EFCAB9*, *GCA*, *PDCD6* scored higher by this metric than the competitors as it highlighted the specific conserved penta-EF hand domain shared by these proteins, rather than the broader categorization of calcium binding (g:GOSt: “calcium ion binding”, Enrichr: “Calcium Ion Binding”, “Metal Ion Binding”, “PERTURB 8787 KO HNF6…” and 27 others). Additionally, we leveraged the MedCPT article encoder to quantify redundancy in top terms. Specifically, we calculated the number of term pairs with high semantic similarity (cosine similarity of embeddings >0.95) amongst a given model’s top terms. By this definition, Enrichr’s aforementioned terms contained the redundant pair “Calcium Ion Binding” and “Metal Ion Binding.”

We then tested whether the models recovered relationships in gene sets designed to represent biological signals. For this task, we utilized the Hallmark Collection (31), which contains fifty gene sets representing well-characterized biological states in cancer identified from gene expression patterns across many microarray and RNAseq experiments (Figure S2A). We randomly sampled 200 queries of sizes 10-100 and found that GeneTEA consistently found terms with higher MedCPT Relevance than the competitors, regardless of query length (Figure 3C, Figure S2B-D). We then randomly combined 100 pairs of these sampled queries and found that although the pairs may have contained entirely unrelated sets of genes, GeneTEA still consistently recovered terms with high MedCPT Relevance (Figure 3C, Figure S2B-D).

Next, we tested the models on experimentally derived queries where a biological relationship was expected but uncertain (Figure 3D, Figure S2E). This included 188 sets of structurally similar proteins identified by clustering AlphaFold2 predictions (32); 152 sets of interacting proteins identified using proximity-dependent biotinylation (BioID) (34); 90 sets of genes with rare variants tied to various diseases by genome-wide association studies (GWAS) across the UK BioBank (35); and finally, 38 expression modules and 64 perturbation clusters where CRISPRi knockdowns produced similar phenotypes in a single-cell Perturb-seq dataset in immortalized cancer cell lines (36). Again, GeneTEA consistently outperformed other models in identifying terms with higher MedCPT Relevance (Figure 3D, Figure S2F-H). When considering the rank order of terms outputted by the models, it was apparent that GeneTEA was better at surfacing the terms with the highest MedCPT Relevance (Figure 3E). Furthermore, GeneTEA’s top terms had lower redundancy than g:GOSt and Enrichr, especially when utilizing the term grouping strategy (Figure 3F). While g:GOSt and Enrichr produced more than 10 redundant term pairs in 77.4% and 93.2% of queries respectively, GeneTEA did so in only 65.6% of queries without term grouping and 36.3% with term grouping.

One example of a structurally similar protein cluster identified with AlphaFold2 - *FCRL2*, *KIR3DS1*, *LILRA6*, and *LILRB3* - highlighted GeneTEA’s ability to identify the biological signal underlying a query (Figure 4A). The top term identified by GeneTEA indicated these genes encode immunoglobulin-like receptors that contain immunoreceptor tyrosine-based inhibitory motifs (ITIM), which are critical for binding MHC class I molecules (37). As this query was constructed by assessing protein structural similarity, it was not surprising that the GeneTEA enrichment emphasized this shared domain. Additionally, the “∼ LRC” and “∼ cluster” terms captured that 3 of the 4 genes encode members of the leukocyte receptor complex (LRC) cluster present on the 19q13 cytoband. The competitor models failed to detect the conserved motif and LRC cluster membership, only finding that *LILRA6* and *LILRB3* are involved in B cell receptor signaling.

**Figure 4.**
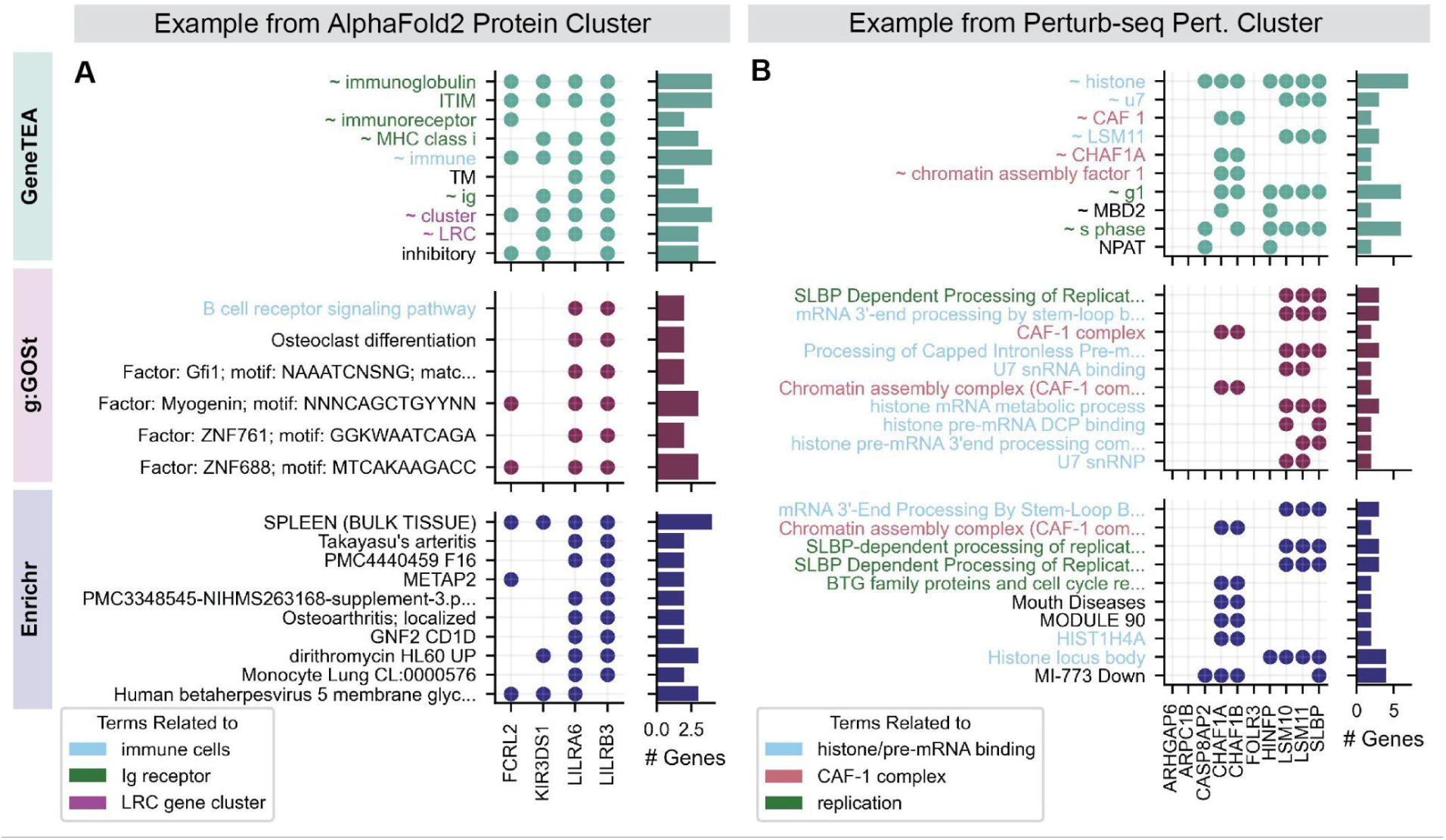
Examples of the top 10 enriched terms from each model. Points in the scatter indicate whether a gene on the x-axis belongs to the gene set described by each term on the y-axis, with number of genes a term appears in on the bar plots to the right. (**A**) Top terms for a set of structurally similar proteins identified from AlphaFold2. (**B**) Top terms for a cluster of Perturb-seq perturbations inducing similar transcriptional phenotypes.

Another example that emphasized the utility of GeneTEA as a hypothesis-building tool was a cluster of perturbations identified in the Perturb-seq dataset (Figure 4B). All three models identify that Chromatin Assembly Factors I (CAF-1) genes *CHAF1A*/*CHAF1B* and *SLBP*/*LSM10*/*LSM11* interact with histones during the cell cycle - the former are histone chaperones that enable nucleosome assembly for DNA replication (38) and the latter are involved in processing replication-dependent histone mRNA (39). However, only GeneTEA identified that *CASP8AP2* and *HINFP* also have replication-related histone function - *CASP8AP2* was shown to regulate the expression of replication-dependent histone mRNA during S phase in colon cancer (40), while *HINFP* is a transcriptional activator that promotes the expression of histone H4 genes at the G1/S phase transition (41). This bolsters a “guilt by association” hypothesis that *CASP8AP2* and *HINFP* are functionally related to CAF-1 and *SLBP* as they produce a similar transcriptional phenotype following knockdown in this Perturb-seq dataset.

### Underlying GeneTEA is a flexible and extensible framework

While GeneTEA was initially established to perform ORA on human genes, its framework could be repurposed to identify enrichment across any entity for which a reasonably large corpus of descriptions can be obtained. As a proof-of-concept, we trained GeneTEA for the budding yeast (*Saccharomyces cerevisiae*) genome utilizing the text from the Alliance of Genome Resources and UniProt (Figure 5A, Supplementary Note S8). We compared the GeneTEA-human and GeneTEA-yeast *tf-idf* matrices under the assumption that orthologous genes should have similar embeddings. To do so, we trained a Nearest Neighbors (NN) model on the GeneTEA-human *tf-idf* matrix and then queried it with yeast ortholog’s *tf-idf* embeddings (Figure 5B, Supplementary Note S9). This NN model had access to all overlapping terms, except for any term representing a protein-coding gene symbol, as this could achieve high performance trivially. For 212 high-confidence human-yeast orthologs identified by the Alliance of Genome Resources, 47.6% of yeast genes had a human ortholog as their nearest neighbor (*k*=1), and 95.8% had a human ortholog within their fifty nearest neighbors (*k*=50). These results demonstrated that GeneTEA could recover shared signals between evolutionarily conserved genes despite being trained on species-specific corpora.

**Figure 5.**
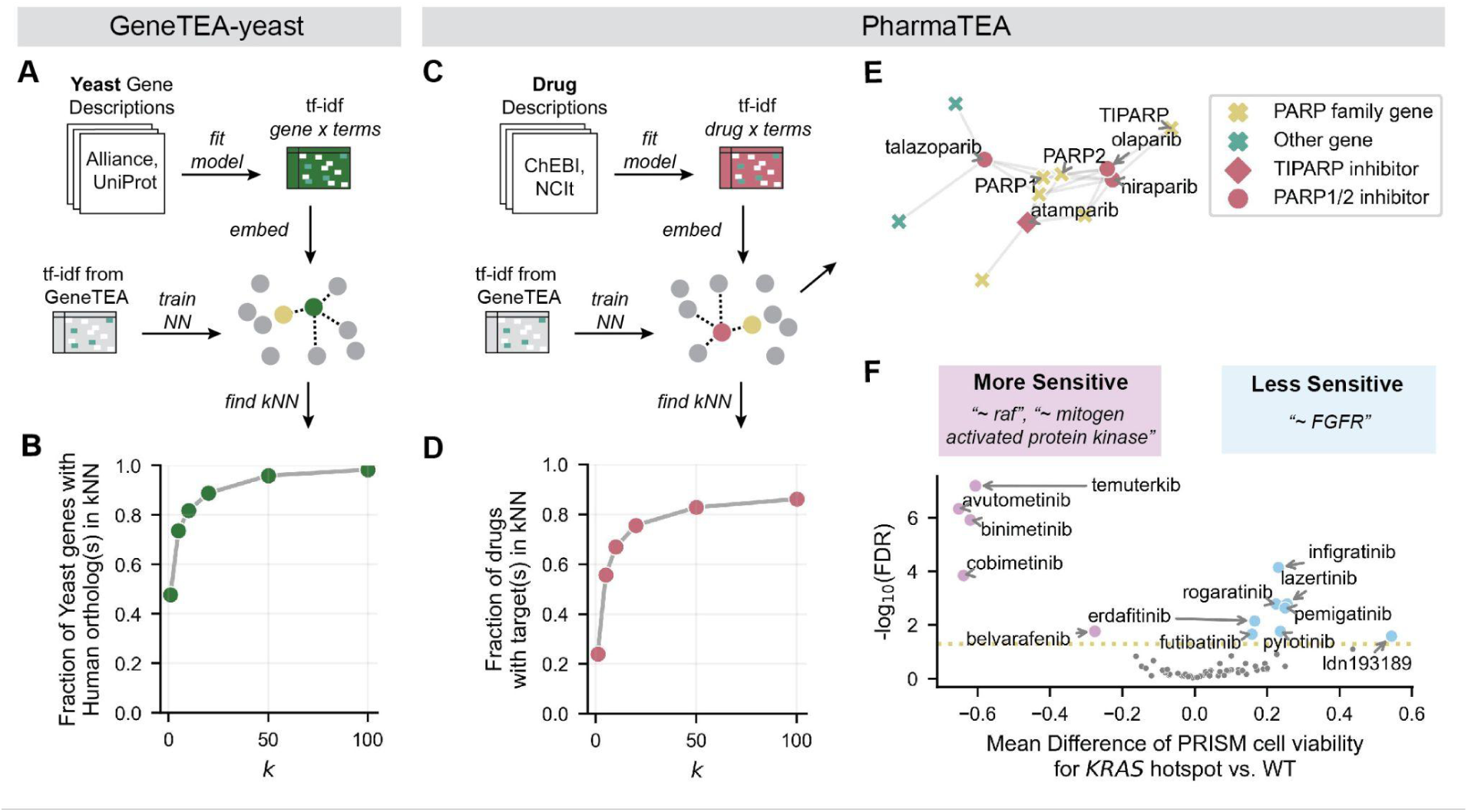
GeneTEA’s flexible framework generalizes to other domains. (**A**) Schema for training GeneTEA-yeast from a corpus of yeast gene descriptions, then finding its *k* nearest neighbors (kNN) GeneTEA (human) genes. (**B**) Fraction of human orthologs recovered in kNN for yeast genes (*n=*212) at various *k* values. (**C**) Schema for training PharmaTEA from a corpus of drug descriptions, finding its kNN GeneTEA genes. (**D**) Fraction of target genes recovered in *k* nearest neighbors for PRISM drugs (*n=*151) at various *k* values. (**E**) Spring layout of a graph with cosine distance weighted edges connecting 5 nearest neighbor genes to PARP inhibitors. (**F**) Top terms for drugs (*n*=64) that *KRAS* mutant cell lines (*n*=159) are more or less sensitive to than *KRAS* WT cell lines (*n*=749), based on two-tailed Student’s *t*-test of the difference in cell viability in the PRISM repurposing dataset.

We further hypothesized that the GeneTEA framework could extend to biological entities other than genes, for instance, small molecules or therapeutic compounds; therefore we obtained drug descriptions from Chemical Entities of Biological Interest (ChEBI) (42) and the National Cancer Institute Thesaurus (NCIt) (43) and trained a new model, PharmaTEA (Figure 5C, Supplementary Note S8). For 151 drugs annotated with PubChemCIDs and genetic targets in the Cancer Dependency Map (DepMap) portal, we repeated the NN analysis comparing GeneTEA to PharmaTEA under the assumption that a drug’s nearest neighbors should contain its annotated genetic target(s). While 23.8% had an annotated target as the nearest neighbor (*k*=1), this improved to 55.6% for the 5 nearest neighbors (*k*=5) (Figure 5D). For example, poly-ADP ribose polymerase (PARP) inhibitors are connected to genes in the PARP family in the 5-NN graph (Figure 5E).

One use case for PharmaTEA was interpreting sensitivity profiles generated in the PRISM repurposing project, for which a variety of FDA-approved compounds were screened across pooled cancer cell lines in an attempt to identify anti-cancer activity (44). When comparing the viability of 159 cell lines with *KRAS* hotspot mutations to 749 wild-type cell lines across 64 drugs in the PRISM repurposing project, PharmaTEA revealed that those harboring mutations are more sensitive to drugs targeting the RAF family and mitogen-activated protein kinases (MAPKs) than the fibroblast growth factor receptor (FGFR) family (Figure 5F, Supplementary Note S10). Oncogenic *KRAS* is upstream of the RAF family and MAPKs, which explains why inhibition of these targets leads to a viability effect. In contrast, *KRAS* and growth factor receptor mutations like *FGFR* and *EGFR* are generally mutually exclusive in cancer, resulting in a lack of sensitivity to the latter in the *KRAS* hotspot population (45).

## DISCUSSION

Identifying the biological signal underlying a list of genes is a common task; however, the success of ORA hinges upon the queried gene set database. With GeneTEA, we demonstrated that a *de novo* gene set database could be learned from free-text gene descriptions using NLP to reduce redundancy while retaining specificity and interpretability. While traditional gene set databases require extensive manual curation, GeneTEA is designed to be updated and expanded automatically. This methodology enables the integration of multiple biological knowledge bases, producing harmonized results with reliable statistics. Furthermore, we showed that GeneTEA’s framework is extensible to other organisms’ genomes or other biological entities such as drugs, so long as a corpus of reliable descriptions can be obtained.

GeneTEA produced fewer false positives while consistently surfacing more relevant enriched terms versus competitor ORA tools. This is likely due to the compact, less overlapping, and more homogeneous nature of the gene sets database in GeneTEA. Additionally, methodological improvements including proper false discovery control, ranking terms based on information content, and filtering redundant terms offered additional benefits over competitors. These results demonstrated that GeneTEA yields reliable and specific enrichment results that enable rapid hypothesis generation.

Furthermore, GeneTEA’s *tf-idf* matrix encoded known relationships between genes. Future work could explore using this embedding in semi-supervised computational biology analyses where prior knowledge of gene function would be beneficial - for example when defining gene programs from expression data or for feature selection in predictive models.

## DATA AVAILABILITY

The data supporting the conclusions of this article are available on Figshare https://doi.org/10.6084/m9.figshare.28635317. GeneTEA’s open-source Python codebase can be found on GitHub at https://github.com/broadinstitute/GeneTEA, with the version used in this article archived on Figshare https://doi.org/10.6084/m9.figshare.28688009.

## AUTHOR CONTRIBUTIONS

Isabella A Boyle: Conceptualization, Methodology, Data curation, Formal analysis, Validation, Software, Visualization, Writing - original draft. Joshua M Dempster: Conceptualization, Supervision, Writing - review and editing. Catarina D Campbell: Supervision, Writing - review and editing. Nayeem Akram Aquib: Software, Writing - review and editing. Mustafa Kocak: Validation, Writing - review and editing. Randy Cresi: Software, Writing - review and editing. Phil G Montgomery: Software, Writing - review and editing.

## SUPPLEMENTARY DATA

*Supplementary Note S1: Human gene description corpus*

To assemble our corpus, we obtained free-text Human gene descriptions from the following sources:

- Alliance: Downloaded *Homo sapiens* gene descriptions from version 8.0.0 (Accessed 2025-02-21) (23)
- CIViC: Downloaded Features TSV from 01-Feb-2025 Release (Accessed 2025-02-21) (22)
- NCBI: Obtained RefSeq summaries from https://ftp.ncbi.nlm.nih.gov/gene/DATA/gene_summary.gz, as well as names/aliases and gene locations in https://ftp.ncbi.nlm.nih.gov/gene/DATA/GENE_INFO/Mammalia/Homo_sapiens.g ene_info.gz (Accessed 2025-02-21) (20)
- UniProt: Downloaded Function, Domain, Subunit structure, etc. sections for all Reviewed (Swiss-prot) Human proteins (Accessed 2025-02-21) (21)

*Supplementary Note S2: Tokenization and phrase extraction*

All gene descriptions in the corpus were mapped to a consistent set of gene symbols using the HGNC symbol mapping (33). Texts were tokenized into sentences using nltk’s sent_tokenize (v3.7) and into words by splitting on spaces. Tokens were converted to lowercase if only one letter was capitalized, otherwise their original case was retained. Candidate phrases were identified using the UMLS2024AB MRCONSO file (accessed 2025-02-24) (24), where English phrases for each concept defined by at least 2 sources were extracted from sentences.

*Supplementary Note S3: Synonym set definition*

Each token is embedded with SapBERT via the huggingface API (transformers v4.46.3) based on code adapted from Hu et al. 2023 (13). First, the AutoTokenizer encoded the input into tokens which were then passed to the AutoModel, followed by mean pooling using the attention mask. These embeddings were aggregated into a matrix, and then sklearn’s HDBSCAN (v1.4.0) was run with min_sample_size=2 and min_cluster_size=2 on the embeddings. All clusters extracted were treated as synonym sets named by the most frequent member of the set with the prefix “∼”.

We compared the learned synonym sets to lists of concept names identified in the English portion of the UMLS2024AB MRCONSO file (accessed 2025-02-24) (24). Concretely, we applied the same tokenization used in GeneTEA to the concept names in the STR column of the file and retained only those overlapping with the GeneTEA vocabulary. We then grouped these concept names by the concept unique identifier (CUI) and retained the 18, 780 CUIs containing at least 2 concept names. Finally, we found 12, 318 GeneTEA synonyms with at least 2 tokens overlapping the UMLS concept names and computed the homogeneity, completeness, and V-measure clustering scores for these 89, 830 tokens with sklearn’s homogeneity_completeness_v_measure function (v1.4.0).

*Supplementary Note S4: Embedding with tf-idf*

We fit an sklearn Tfidfvectorizer (v1.4.0) on the tokenized documents, with the following hyperparameters: max_df=0.6, min_df=3, binary=False.

*Supplementary Note S5: Statistical test and effect sizes for overrepresentation*

Overrepresentation in GeneTEA is measured using a hypergeometric test. The scipy.stats (46) implementation hypergeom.sf with loc=1 was used to obtain *p*-values, which were then converted to FDR values using Benjamini-Hochberg’s false discovery control (scipy.stats.false_discovery_control). Version 1.11.1 of scipy.stats was used for this computation. The reported “Effect Size” is the sum of *tf-idf* values per term across the query genes. Additionally, the sum of *tf-idf* values across all genes in the matrix is reported as the “Total Info.”

*Supplementary Note S6: Custom stopwords*

We found that uninteresting, non-biological terms occasionally appeared enriched - such as “exhibits” or “several” - and used the following strategies to exclude them. First, we took the average SapBERT embedding of the top 500 most frequent words in the 2024 MEDLINE 1-gram set (47) and added any terms in the vocabulary with a high Pearson correlation (>0.65). Then, we added phrases made up entirely of stopwords. From this list of stopwords, we identified synonym sets made up of >50% stopwords. Finally, we added terms that appeared in >15% of genes. Any fully uppercase terms or those matching the chromosomal location patterns were not included in this list of custom stopwords.

*Supplementary Note S7: Graph-based filtering and grouping strategy*

The relationship between query genes and enriched terms can naturally be represented as a bipartite graph, where top nodes are genes and bottom nodes are terms with un-weighted edges representing the presence of a term in a gene’s description. Therefore, we leveraged existing implementations of several graph algorithms from the Python package networkx (v3.2.1) (48) to filter out strictly redundant terms and group conceptually similar terms.

First, a bottom node projection of terms is created where edges are present if a pair of terms is present in a fully overlapping set of genes. Then, edges are pruned such that only sub-terms (i.e. “olfactory” and “olfactory receptor”) remain connected. All terms are then ranked by “Effect Size”, “FDR”, “Total Info”, and finally lexicographically. Finally, term nodes are pruned if the term is not the highest ranked amongst its connected nodes, resulting in a filtered set of terms.

Next, terms are grouped based on the intuition that terms with similar meanings will have similar *tf-idf* embeddings within a query and across all genes. Therefore, a new graph is created where nodes are filtered terms and edges have weight

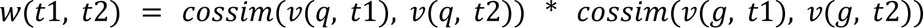

where *v* is the *tf-idf* vector over query genes *q* or all genes *g* for terms *t1, t2* and *cossim* refers to cosine similarity. If *w* < 0.15, the edge is removed to prevent collapsing largely unrelated terms. Then, groups of terms are identified using the networkx greedy_modular_communities function and are labeled by the three highest-ranked terms.

*Supplementary Note S8: Training GeneTEA-yeast and PharmaTEA*

Free-text yeast gene descriptions for GeneTEA-yeast were obtained from the following sources:

- Alliance: Downloaded *Saccharomyces cerevisiae* gene descriptions from version 8.0.0 (Accessed 2025-02-24) (23)
- UniProt: Downloaded Function, Domain, Subunit structure, etc. sections for all Reviewed (Swiss-prot) *S. cerevisiae* proteins (Accessed 2025-02-24) (21)

Free-text compound descriptions for PharmaTEA were obtained from the following sources:

- ChEBI: Downloaded Record Descriptions (Compound) from PubChem PUG View API (Accessed 2025-02-25) (49)
- NCIt: Downloaded Record Descriptions (Compound) from PubChem PUG View API (Accessed 2025-02-25) (49)

Both models were trained identically to GeneTEA, except that GeneTEA’s synonym sets were used to ensure alignment of term definitions.

*Supplementary Note S9: k Nearest Neighbors analysis*

To compare GeneTEA-yeast and PharmaTEA embeddings to GeneTEA, we ran a *k* Nearest Neighbor (kNN) analysis. First, overlapping terms were identified; though terms that matched the protein-coding gene symbols were excluded as they could trivially connect neighbors. The *tf-idf* embedding from GeneTEA was then used to train an sklearn NearestNeighbors model (v1.4.0) with metric=“cosine”. The kNN of each yeast ortholog or drug were then obtained for *k*=1, 5, 10, 20, 50, and 100, and the number of human orthologs/genetic targets at each *k* value was counted. Yeast-human orthologs were obtained from Alliance of Genome Resources (version 8.0.0, accessed 2025-02-21) “Alliance combined orthology data” TSV (23), filtered to only ortholog pairs identified by all 9 algorithms. Drug-target annotations were obtained from the PortalCompounds.csv in the DepMap 24Q4 Public Figshare (50).

*Supplementary Note S10: PRISM repurposing sensitivity for KRAS mutant vs. wild-type cell lines*

We obtained the PRISM repurposing log2-fold change data, where cell viability was measured as the difference in PRISM barcode abundance between treatment and DMSO conditions, from the file Repurposing_Public_24Q2_Extended_Primary_Data_Matrix.csv in the Repurposing Public 24Q2 Figshare (51). Using hotspot mutation calls from the OmicsSomaticMutationsMatrixHotspot.csv file on the DepMap 24Q4 Public Figshare (50), we performed a two-tailed Student’s *t*-test (scipy.stats.ttest_ind) between the populations with and without a *KRAS* mutation for the 137 drugs where PubMed CIDs and targets had been annotated. We applied Benjimini-Hochberg false discovery control (scipy.stats.false_discovery_control), then called significant compounds based on an FDR threshold of 0.05, with directionality assigned based on the mean difference between the *KRAS* hotspot and WT cell lines. Lists of significantly more sensitive and less sensitive drugs were used to query PharmaTEA, where the background was specified as the 137 drugs on which the *t*-test was performed. Version 1.11.1 of scipy.stats was used throughout this analysis.

---

**Supplementary Figure S1.**
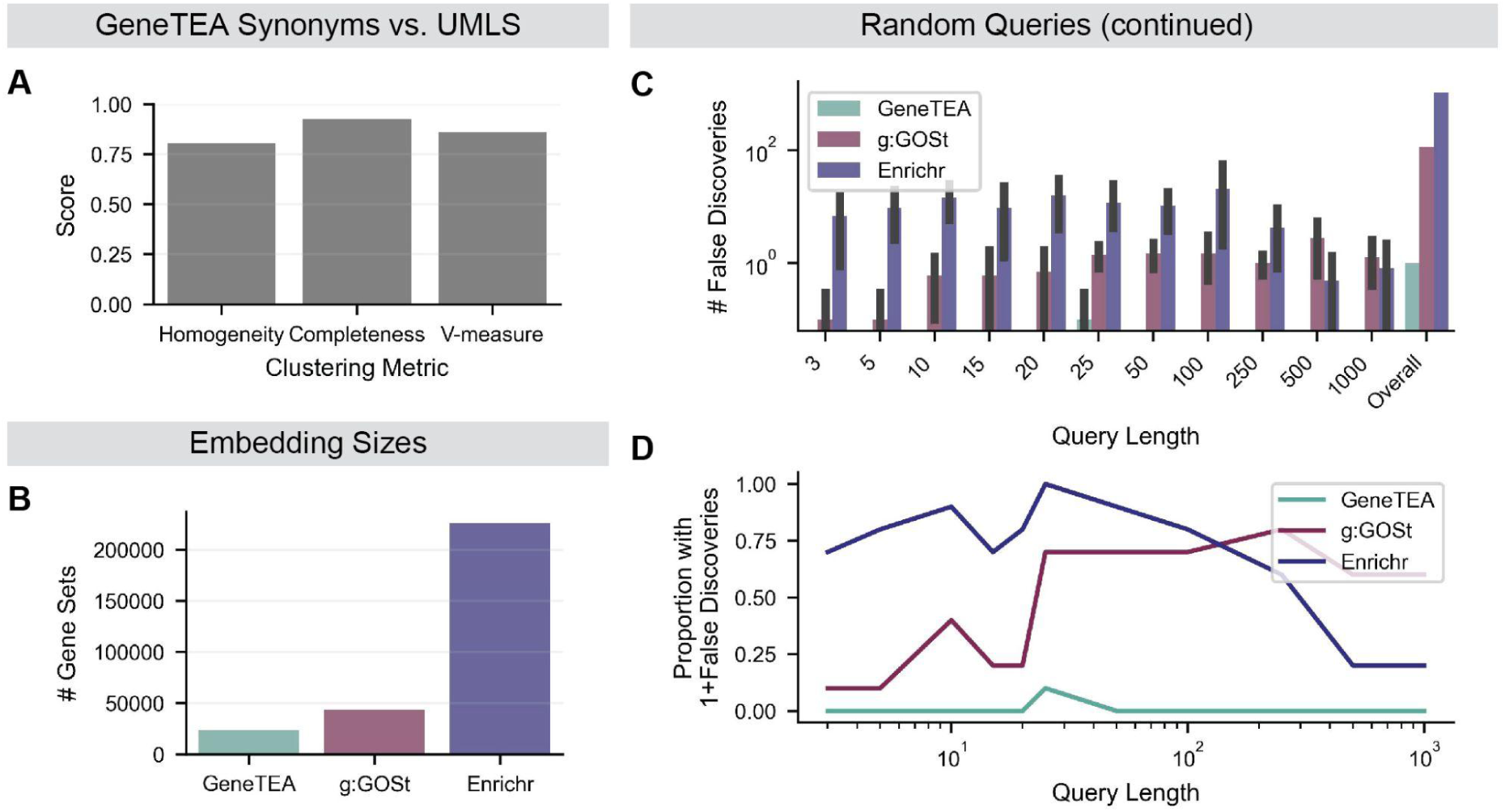
Additional information on embedding construction and false discovery control. (**A**) Scores from various clustering metrics when comparing GeneTEA’s synonym sets to UMLS Metathesaurus concepts (*n*=89,830 tokens). (**B**) Number of gene sets in each model’s embedding. (**C**) Bar plot showing the number of false discoveries for all lengths of queries and overall, with error bars representing the standard error of the mean over 10 samples. (**D**) Proportion of random queries with false discoveries versus length of query.

**Supplementary Figure S2.**
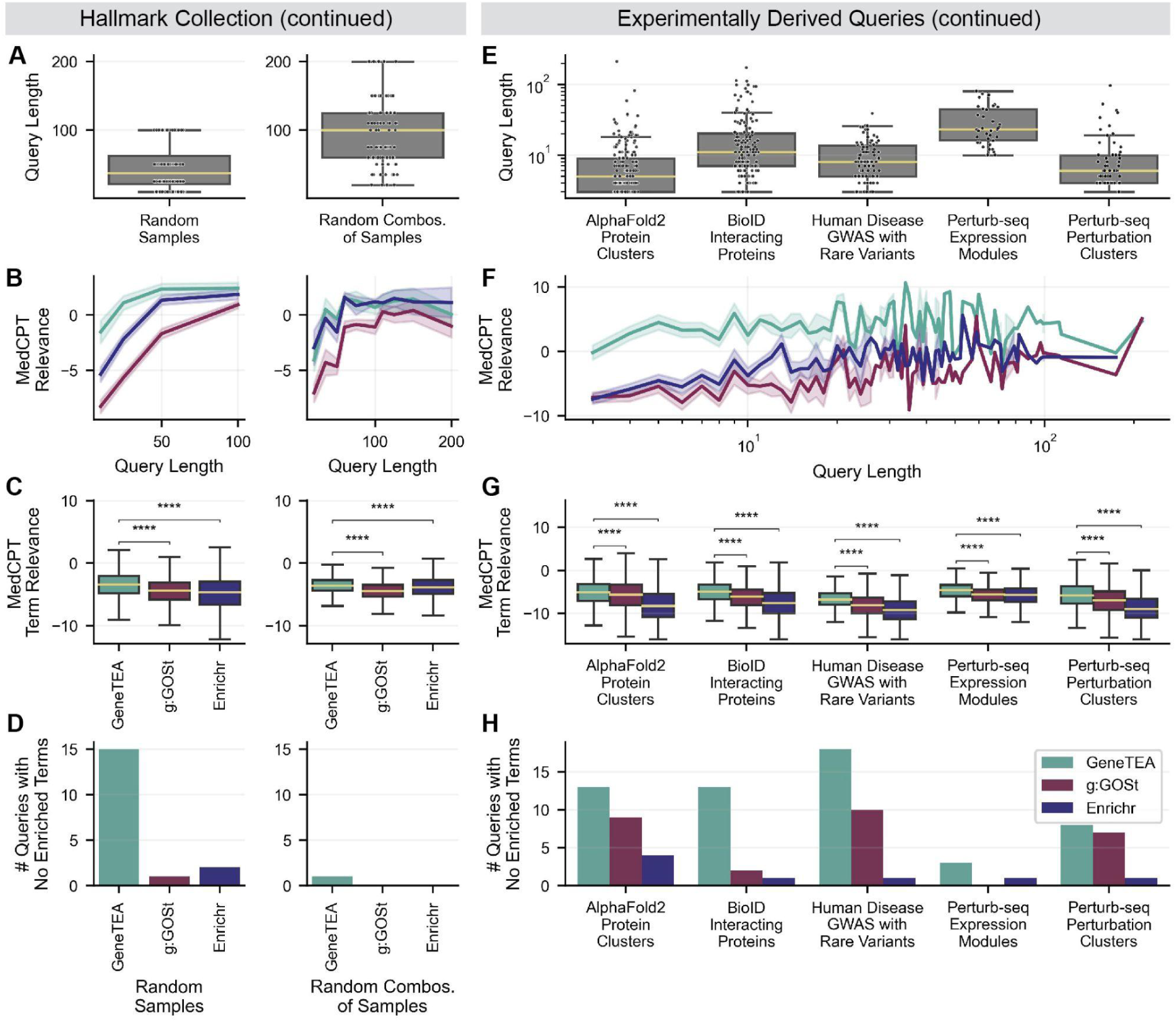
Additional information on benchmarking tests. (**A**) Lengths of queries in Hallmark Collection (*n*=200 in left, *n*=100 in right). (**B**) MedCPT Relevance versus length of query in the Hallmark Collection, with error bars representing the standard error of the mean. (**C**) MedCPT Relevance for each Hallmark Collection query’s top 100 enriched terms. (**D**) Number of queries where no enriched terms were detected in the Hallmark Collection. (**E**) Lengths of queries in experimentally derived queries (*n*=532). (**F**) MedCPT Relevance versus length of query in experimentally derived queries, with error bars representing the standard error of the mean. (**G**) MedCPT Relevance for each experimentally derived query’s top 100 enriched terms. (**H**) Number of experimentally derived queries where no enriched terms were detected in experimentally derived queries. All significance values in **C** and **G** are from a right-tailed Student’s *t*-test, where **** *p*-value < 0.0001.

## REFERENCES

1. Das, S., McClain, C.J. and Rai, S.N. (2020) Fifteen Years of Gene Set Analysis for High-Throughput Genomic Data: A Review of Statistical Approaches and Future Challenges. Entropy, 22.

2. Harris, M.A., Clark, J., Ireland, A., Lomax, J., Ashburner, M., Foulger, R., Eilbeck, K., Lewis, S., Marshall, B., Mungall, C., et al. (2004) The Gene Ontology (GO) database and informatics resource. Nucleic Acids Res., 32, D258–61.

3. Robinson, P.N., Köhler, S., Bauer, S., Seelow, D., Horn, D. and Mundlos, S. (2008) The Human Phenotype Ontology: a tool for annotating and analyzing human hereditary disease. Am. J. Hum. Genet., 83, 610–615.

4. Subramanian, A., Tamayo, P., Mootha, V.K., Mukherjee, S., Ebert, B.L., Gillette, M.A., Paulovich, A., Pomeroy, S.L., Golub, T.R., Lander, E.S., et al. (2005) Gene set enrichment analysis: a knowledge-based approach for interpreting genome-wide expression profiles. Proc. Natl. Acad. Sci. U. S. A., 102, 15545–15550.

5. Ogata, H., Goto, S., Sato, K., Fujibuchi, W., Bono, H. and Kanehisa, M. (1999) KEGG: Kyoto Encyclopedia of Genes and Genomes. Nucleic Acids Res., 27, 29–34.

6. Pico, A.R., Kelder, T., van Iersel, M.P., Hanspers, K., Conklin, B.R. and Evelo, C. (2008) WikiPathways: pathway editing for the people. PLoS Biol., 6, e184.

7. Joshi-Tope, G., Gillespie, M., Vastrik, I., D’Eustachio, P., Schmidt, E., de Bono, B., Jassal, B., Gopinath, G.R., Wu, G.R., Matthews, L., et al. (2005) Reactome: a knowledgebase of biological pathways. Nucleic Acids Res., 33, D428–32.

8. Kolberg, L., Raudvere, U., Kuzmin, I., Adler, P., Vilo, J. and Peterson, H. (2023) g:Profiler-interoperable web service for functional enrichment analysis and gene identifier mapping (2023 update). Nucleic Acids Res., 51, W207–W212.

9. Xie, Z., Bailey, A., Kuleshov, M.V., Clarke, D.J.B., Evangelista, J.E., Jenkins, S.L., Lachmann, A., Wojciechowicz, M.L., Kropiwnicki, E., Jagodnik, K.M., et al. (2021) Gene Set Knowledge Discovery with Enrichr. Curr Protoc, 1, e90.

10. Maleki, F. and Kusalik, A.J. (2020) Gene Set Overlap: An Impediment to Achieving High Specificity in Over-representation Analysis. bioRxiv, 10.1101/319145.

11. Karp, P.D., Midford, P.E., Caspi, R. and Khodursky, A. (2021) Pathway size matters: the influence of pathway granularity on over-representation (enrichment analysis) statistics. BMC Genomics, 22, 191.

12. Galke, L., Diera, A., Lin, B.X., Khera, B., Meuser, T., Singhal, T., Karl, F. and Scherp, A. (2022) Are We Really Making Much Progress in Text Classification? A Comparative Review. arXiv [cs.CL].

13. Hu, M., Alkhairy, S., Lee, I., Pillich, R.T., Bachelder, R., Ideker, T. and Pratt, D. (2023) Evaluation of large language models for discovery of gene set function. Res Sq, 10.21203/rs.3.rs-3270331/v1.

14. Joachimiak, M.P., Caufield, J.H., Harris, N.L., Kim, H. and Mungall, C.J. (2023) Gene Set Summarization using Large Language Models. ArXiv, 10.1093/database/baw110.

15. Wang, Z., Jin, Q., Wei, C.-H., Tian, S., Lai, P.-T., Zhu, Q., Day, C.-P., Ross, C. and Lu, Z. (2024) GeneAgent: Self-verification language agent for gene set knowledge discovery using domain databases. ArXiv.

16. Chen, S., Li, Y., Lu, S., Van, H., Aerts, H.J.W.L., Savova, G.K. and Bitterman, D.S. (2024) Evaluating the ChatGPT family of models for biomedical reasoning and classification. J. Am. Med. Inform. Assoc., 31, 940–948.

17. Valentini, M., Szkandera, J., Smolle, M., Scheipl, S., Leithner, A. and Andreou, D. (2024) Artificial intelligence large language model ChatGPT: is it a trustworthy and reliable source of information for sarcoma patients? Front Public Health, 12, 1303319.

18. Morgan, S.L., Naderi, P., Koler, K., Pita-Juarez, Y., Prokopenko, D., Vlachos, I.S., Tanzi, R.E., Bertram, L. and Hide, W.A. (2022) Most Pathways Can Be Related to the Pathogenesis of Alzheimer’s Disease. Front. Aging Neurosci., 14, 846902.

19. Leong, H.S. and Kipling, D. (2009) Text-based over-representation analysis of microarray gene lists with annotation bias. Nucleic Acids Res., 37, e79.

20. O’Leary, N.A., Wright, M.W., Brister, J.R., Ciufo, S., Haddad, D., McVeigh, R., Rajput, B., Robbertse, B., Smith-White, B., Ako-Adjei, D., et al. (2016) Reference sequence (RefSeq) database at NCBI: current status, taxonomic expansion, and functional annotation. Nucleic Acids Res., 44, D733–45.

21. UniProt Consortium (2023) UniProt: the Universal Protein Knowledgebase in 2023. Nucleic Acids Res., 51, D523–D531.

22. Krysiak, K., Danos, A.M., Saliba, J., McMichael, J.F., Coffman, A.C., Kiwala, S., Barnell, E.K., Sheta, L., Grisdale, C.J., Kujan, L., et al. (2023) CIViCdb 2022: evolution of an open-access cancer variant interpretation knowledgebase. Nucleic Acids Res., 51, D1230–D1241.

23. Bult, C.J. and Sternberg, P.W. (2023) The alliance of genome resources: transforming comparative genomics. Mamm. Genome, 34, 531–544.

24. Bodenreider, O. (2004) The Unified Medical Language System (UMLS): integrating biomedical terminology. Nucleic Acids Res., 32, D267–70.

25. Liu, F., Shareghi, E., Meng, Z., Basaldella, M. and Collier, N. (2020) Self-Alignment Pretraining for Biomedical Entity Representations. arXiv [cs.CL].

26. Campello, R.J.G.B., Moulavi, D. and Sander, J. (2013) Density-based clustering based on hierarchical density estimates. In Advances in Knowledge Discovery and Data Mining, Lecture notes in computer science. Springer Berlin Heidelberg, Berlin, Heidelberg, pp. 160–172.

27. Aizawa, A. (2003) An information-theoretic perspective of tf–idf measures. Inf. Process. Manag., 39, 45–65.

28. Razzaque, M.A., Nishizawa, T., Komoike, Y., Yagi, H., Furutani, M., Amo, R., Kamisago, M., Momma, K., Katayama, H., Nakagawa, M., et al. (2007) Germline gain-of-function mutations in RAF1 cause Noonan syndrome. Nat. Genet., 39, 1013–1017.

29. Pedregosa, F., Varoquaux, G., Gramfort, A., Michel, V., Thirion, B., Grisel, O., Blondel, M., Louppe, G., Prettenhofer, P., Weiss, R., et al. (2011) Scikit-learn: Machine Learning in Python. J. Mach. Learn. Res., **abs/**1201.0490.

30. Jin, Q., Kim, W., Chen, Q., Comeau, D.C., Yeganova, L., Wilbur, W.J. and Lu, Z. (2023) MedCPT: Contrastive Pre-trained Transformers with large-scale PubMed search logs for zero-shot biomedical information retrieval. Bioinformatics, 39, btad651.

31. Liberzon, A., Birger, C., Thorvaldsdóttir, H., Ghandi, M., Mesirov, J.P. and Tamayo, P. (2015) The Molecular Signatures Database (MSigDB) hallmark gene set collection. Cell Syst., 1, 417–425.

32. Barrio-Hernandez, I., Yeo, J., Jänes, J., Mirdita, M., Gilchrist, C.L.M., Wein, T., Varadi, M., Velankar, S., Beltrao, P. and Steinegger, M. (2023) Clustering predicted structures at the scale of the known protein universe. Nature, 622, 637–645.

33. Seal, R.L., Braschi, B., Gray, K., Jones, T.E.M., Tweedie, S., Haim-Vilmovsky, L. and Bruford, E.A. (2023) Genenames.org: the HGNC resources in 2023. Nucleic Acids Res., 51, D1003–D1009.

34. Go, C.D., Knight, J.D.R., Rajasekharan, A., Rathod, B., Hesketh, G.G., Abe, K.T., Youn, J.-Y., Samavarchi-Tehrani, P., Zhang, H., Zhu, L.Y., et al. (2021) A proximity-dependent biotinylation map of a human cell. Nature, 595, 120–124.

35. Wang, Q., Dhindsa, R.S., Carss, K., Harper, A.R., Nag, A., Tachmazidou, I., Vitsios, D., Deevi, S.V.V., Mackay, A., Muthas, D., et al. (2021) Rare variant contribution to human disease in 281, 104 UK Biobank exomes. Nature, 597, 527–532.

36. Replogle, J.M., Saunders, R.A., Pogson, A.N., Hussmann, J.A., Lenail, A., Guna, A., Mascibroda, L., Wagner, E.J., Adelman, K., Lithwick-Yanai, G., et al. (2022) Mapping information-rich genotype-phenotype landscapes with genome-scale Perturb-seq. Cell, 185, 2559–2575.e28.

37. Bern, M.D., Beckman, D.L., Ebihara, T., Taffner, S.M., Poursine-Laurent, J., White, J.M. and Yokoyama, W.M. (2017) Immunoreceptor tyrosine-based inhibitory motif-dependent functions of an MHC class I-specific NK cell receptor. Proc. Natl. Acad. Sci. U. S. A., 114, E8440–E8447.

38. Franklin, R., Guo, Y., He, S., Chen, M., Ji, F., Zhou, X., Frankhouser, D., Do, B.T., Chiem, C., Jang, M., et al. (2022) Regulation of chromatin accessibility by the histone chaperone CAF-1 sustains lineage fidelity. Nat. Commun., 13, 2350.

39. Armstrong, C. and Spencer, S.L. (2021) Replication-dependent histone biosynthesis is coupled to cell-cycle commitment. Proc. Natl. Acad. Sci. U. S. A., 118, e2100178118.

40. Hummon, A.B., Pitt, J.J., Camps, J., Emons, G., Skube, S.B., Huppi, K., Jones, T.L., Beissbarth, T., Kramer, F., Grade, M., et al. (2012) Systems-wide RNAi analysis of CASP8AP2/FLASH shows transcriptional deregulation of the replication-dependent histone genes and extensive effects on the transcriptome of colorectal cancer cells. Mol. Cancer, 11, 1.

41. Medina, R., van Wijnen, A.J., Stein, G.S. and Stein, J.L. (2006) The histone gene transcription factor HiNF-P stabilizes its cell cycle regulatory co-activator p220NPAT. Biochemistry, 45, 15915–15920.

42. Hastings, J., Owen, G., Dekker, A., Ennis, M., Kale, N., Muthukrishnan, V., Turner, S., Swainston, N., Mendes, P. and Steinbeck, C. (2016) ChEBI in 2016: Improved services and an expanding collection of metabolites. Nucleic Acids Res., 44, D1214–9.

43. Fragoso, G., de Coronado, S., Haber, M., Hartel, F. and Wright, L. (2004) Overview and utilization of the NCI thesaurus. Comp. Funct. Genomics, 5, 648–654.

44. Corsello, S.M., Nagari, R.T., Spangler, R.D., Rossen, J., Kocak, M., Bryan, J.G., Humeidi, R., Peck, D., Wu, X., Tang, A.A., et al. (2020) Discovering the anti-cancer potential of non-oncology drugs by systematic viability profiling. *Nat*. Cancer, 1, 235–248.

45. Huang, L., Guo, Z., Wang, F. and Fu, L. (2021) KRAS mutation: from undruggable to druggable in cancer. Signal Transduct. Target. Ther., 6, 386.

46. Virtanen, P., Gommers, R., Oliphant, T.E., Haberland, M., Reddy, T., Cournapeau, D., Burovski, E., Peterson, P., Weckesser, W., Bright, J., et al. (2020) SciPy 1.0: fundamental algorithms for scientific computing in Python. Nat. Methods, 17, 261–272.

47. Lu, C.J., Tormey, D., McCreedy, L. and Browne, A.C. (2015) Generating the MEDLINE N-gram set. Conference of American Medical Informatics Association.

48. Hagberg, A., Swart, P.J. and Schult, D.A. (2008) Exploring network structure, dynamics, and function using NetworkX Los Alamos National Laboratory (LANL), Los Alamos, NM (United States).

49. Kim, S., Chen, J., Cheng, T., Gindulyte, A., He, J., He, S., Li, Q., Shoemaker, B.A., Thiessen, P.A., Yu, B., et al. (2023) PubChem 2023 update. Nucleic Acids Res., 51, D1373–D1380.

50. DepMap, B. (2024) DepMap 24Q4 Public. 10.25452/FIGSHARE.PLUS.27993248.V1.

51. DepMap, B. and Kocak, M. (2024) Repurposing Public 24Q2. 10.6084/M9.FIGSHARE.25917643.V1.

